# Approximation to the distribution of fitness effects across functional categories in human segregating polymorphisms

**DOI:** 10.1101/002345

**Authors:** Fernando Racimo, Joshua G. Schraiber

## Abstract

Quantifying the proportion of polymorphic mutations that are deleterious or neutral is of fundamental importance to our understanding of evolution, disease genetics and the maintenance of variation genome-wide. Here, we develop an approximation to the distribution of fitness effects (DFE) of segregating single-nucleotide mutations in humans. Unlike previous methods, we do not assume that synonymous mutations are neutral or not strongly selected, and we do not rely on fitting the DFE of all new nonsynonymous mutations to a single probability distribution, which is poorly motivated on a biological level. We rely on a previously developed method that utilizes a variety of published annotations (including conservation scores, protein deleteriousness estimates and regulatory data) to score all mutations in the human genome based on how likely they are to be affected by negative selection, controlling for mutation rate. We map this score to a scale of fitness coefficients via maximum likelihood using diffusion theory and a Poisson random field model on SNP data. Our method serves to approximate the deleterious DFE of mutations that are segregating, regardless of their genomic consequence. We can then compare the proportion of mutations that are negatively selected or neutral across various categories, including different types of regulatory sites. We observe that the distribution of intergenic polymorphisms is highly peaked at neutrality, while the distribution of nonsynonymous polymorphisms is bimodal, with a neutral peak and a second peak at *s ≈ −*10*^−^*^4^. Other types of polymorphisms have shapes that fall roughly in between these two. We find that transcriptional start sites, strong CTCF-enriched elements and enhancers are the regulatory categories with the largest proportion of deleterious polymorphisms.

**Author Summary:** The relative frequencies of polymorphic mutations that are deleterious, nearly neutral and neutral is traditionally called the distribution of fitness effects (DFE). Obtaining an accurate approximation to this distribution in humans can help us understand the nature of disease and the mechanisms by which variation is maintained in the genome. Previous methods to approximate this distribution have relied on fitting the DFE of new mutations to a single probability distribution, like a normal or an exponential distribution. Generally, these methods also assume that a particular category of mutations, like synonymous changes, can be assumed to be neutral or nearly neutral. Here, we provide a novel method designed to reflect the strength of negative selection operating on any segregating site in the human genome. We use a maximum likelihood mapping approach to fit these scores to a scale of neutral and negative fitness coefficients. Finally, we compare the shape of the DFEs we obtain from this mapping for different types of functional categories. We observe the distribution of polymorphisms has a strong peak at neutrality, as well as a second peak of deleterious effects when restricting to nonsynonymous polymorphisms.

## Introduction

Genetic variation within species is shaped by a variety of evolutionary processes, including mutation, demography, and natural selection. With the advent of whole-genome sequencing, we can make unprecedented inferences about these and other processes by analyzing population genomic data. An important goal is to understand the extent to which segregating genetic variants are impacted by natural selection, and to quantify the intensity of natural selection acting genome-wide. Understanding the prevalence of different modes of selection on a genomic scale has wide-ranging implications across evolutionary and medical genetics. For instance, genome-wide association studies (GWAS) are searching for mutations associated with disease in large samples of humans [1]. Because mutations associated with disease are *a priori* likely to be deleterious, quantifying the portion of mutations that are deleterious along with their average effects could have significant implications for the design and interpretation of GWAS. Moreover, the ENCODE project has recently claimed that much of the genome is involved in some form of functional activity [2,3]. However, the extent to which molecular signals identified by this project are actually produced by biological processes affecting fitness has been disputed [4, 5]. Indeed, comparative genomics studies suggest that only a small proportion of the human genome (5 *–* 10%) is under purifying selection, based on signals detectable on phylogenetic timescales [6–8]. Quantifying the DFE in noncoding regions is a first step toward understanding the fitness implications of functional activity at the genomic level.

Traditionally, studies have sought to estimate the distribution of fitness effects (DFE) for nonsynonymous mutations by using summary statistics based on the number of polymorphisms and substitutions [9–11] and/or the full frequency spectrum [12–14]. These studies typically assumed that synonymous variation is neutral or under weak selection. Many of these analyses suggest that while a large proportion of nonsynonymous mutations are nearly neutral, there is also a significant probability that an amino acid changing mutation will be strongly deleterious. While these studies were limited to analysis of protein-coding genes, recent work has focused on quantifying the DFE in regulatory regions, including short interspersed genomic elements such as enhancers [15, 16] and cis-regulatory regions [17]. Reviews of these and other approaches can be found in ref. [18, 19].

There are several obstacles to quantifying the DFE of new or segregating mutations genome-wide. First, inferences about the DFE are confounded by demography [20]. For example, a high proportion of low frequency derived alleles is a signature of negative selection, but can also be caused by recent population growth [21]. Hence, a well-supported demographic model must be used to appropriately control for population history when inferring the DFE. Second, most current methods rely on dividing up polymorphisms into either putatively selected sites or putatively neutral (or less selected) sites (for example, nonsynonymous and synonymous sites, respectively). These studies have relied on fitting a demographic moel to the neutral class of sites and then fitting the DFE of new mutations to a probability distribution, typically an exponential or gamma distribution [9,13] to the class of sites that are putatively under selection (e.g. nonsynonymous sites). While flexible, these distributions may miss some important features of the DFE [22]. For example, mutation accumulation experiments suggest that the DFE may be bimodal for at least some species, with most mutations either having nearly neutral or strongly deleterious effects, and very few mutations falling in between [23, 24]. Thus, assuming two classes of sites may not serve to capture all the relevant information about the DFE (but see [25] for an example of fitting a multimodal DFE to population genetic data and [22, 26] for nonparametric approaches to estimating the DFE of new amino-acid changing mutations). Finally, previous studies have been restricted to analyzing specific subclasses of mutations (e.g. nonsynonymous, enhancers, etc.) because until recently, no single metric existed that could serve to compare the disruptive potential of any type of variant, regardless of its genomic consequence.

Recently, Kircher et al. [27] developed a method to synthesize a large number of annotations into a single score to predict the pathogenicity or disruptive potential of any mutation in the genome. It is based on an analysis comparing real and simulated changes that occurred in the human lineage since the human-chimpanzee ancestor, and that are now fixed in present-day humans. The method relies on the realistic assumption that the set of real changes is depleted of deleterious variation due to the action of negative selection, which has pruned away disruptive variants, while the simulated set is not depleted of such variation. A support vector machine (SVM) was trained to distinguish the real from the simulated changes using a kernel of 63 annotations (including conservation scores, regulatory data and protein deleteriousness scores), and then used to assign a score (C-score) to all possible single-nucleotide changes in the human genome, controlling for local variation in mutation rates. These C-scores are meant to be predictors of how disruptive a given change may be, and are comparable across all types of sites (nonsynonymous, synonymous, regulatory, intronic or intergenic). Thus, they allow for a strict ranking of predicted functional disruption for mutations that may not be otherwise comparable. C-scores are PHRED scaled, with larger values corresponding to more disruptive effects.

Importantly, human-specific genetic variation patterns are not used as input to train the C-score SVM. In this work, we make use of the C-scores to provide a fine-grained stratification of the deleteriousness of variants segregating in modern human populations. We take advantage of the Poisson random field model [28, 29] with a realistic model of human demographic history to fit a maximum likelihood selection coefficient for each C-score, creating a mapping from C-scores to selection coefficients.

## Results

### A mapping from C-scores to selection coefficients

To map C-scores to selective coefficients, we obtained allele frequency information from 9 Yoruba (YRI) individuals (18 haploid sequences) sequenced to high-coverage using whole-genome shotgun sequencing as part of a dataset produced by Complete Genomics (CG) [30]. We removed sites that had a Duke Uniqueness 20bp-mapability score *<* 1 (downloaded from the UCSC Genome Browser, [31]), to avoid potential errors due to mismapping or miscalling in regions of the genome that are not uniquely mapable.

When inferring the DFE, we focused only on models of neutral evolution and negative selection, because C-scores are uninformative about adaptive vs. deleterious disruption (i.e. a high C-score could either reflect a highly deleterious change or a highly adaptive change). Additionally, because we are using polymorphism data only, positive selection should contribute little to the site-frequency spectrum [32].

We first binned polymorphisms into C-scores rounded to the nearest integer and computed the site frequency spectrum for each bin (Figure S1). We then fit the lowest possible C-score (C= 0), presumed to be neutral, to different models of demographic history. We computed the likelihood of the SFS in this bin for a constant population size model, a range of exponential growth models, the model inferred by Tennessen et al. [33] and the model inferred by Harris and Nielsen [34] from the distribution of tracts of identity by state (IBS) (Figure S2), and used an EM algorithm to correct for ancestral state misidentification (Figure S3, see Materials and Methods). We find that a model of exponential growth at population-scaled rate = 1 for 13,000 generations is the best fit to the corrected SFS, although the Tennessen model is almost as good a fit (Figure S2).

Using the best-fitting demography, we next fit a range of models with different selection coefficients to the EM-corrected site frequency spectrum for each C-score bin, using maximum likelihood (Figure 1.A) (see Methods). We restricted to C *≤* 40, because very few sites have larger C-scores, and hence estimates of the selection coefficients for those C-scores are unreliable. We tested the robustness of the mappings to different levels of background selection, by partitioning the data into deciles of B-scores [35] and re-computing the C-to-s mapping for each decile. We observe that the mapping is generally robust to background selection, with substantial differences only observed at the lowest two B-score quantiles, which correspond to high background selection (Figure S4). For this reason, and so as to obtain reliable DFEs at exonic sites (where background selection is generally higher than in the rest of the genome), we also performed a neutral demographic fitting and a C-to-s mapping while restricting only to sites in the exome (Figure 1.C). This mapping has a steeper decline than the genomic mapping, reflecting patterns of background selection which are not fully controlled by C-scores but that affects the SFS. We therefore show estimated DFEs using both the genome-wide and the exome-wide fittings below. After removing the C-score bins that best fit the neutral model, we fit a smoothing spline with 20 degrees of freedom to the remaining C-scores, so as to produce a continuous mapping of C-scores to selection coefficients (Figure 1.A).

**Figure 1.**
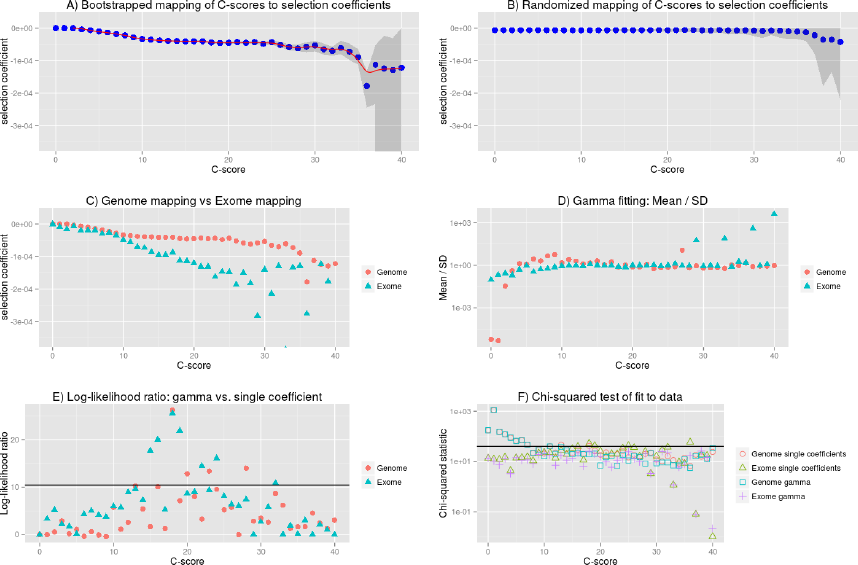
Mapping of C-scores to selection coefficients. A) We first fit a single coefficient to each bin using data from all autosomes in the genome. Red dots represent the maximum likelihood selection coefficient corresponding to each C-score bin. The blue line is a smooth spline fitted to the discrete mappings of C-scores to log-scaled selection coefficients after excluding the neutral bins (degrees of freedom = 20). The grey shade is a 95% confidence interval obtained from bootstrapping the data 100 times in each bin. B) To verify the shape of the mapping was not a result of the number of sites in each bin, we randomized the C-score labels across polymorphisms, and recomputed the mapping. C) To account for exonic patterns of conservation and background selection that may not have been captured by C-scores, we computed a mapping based solely on exonic sites. D) We fitted gamma distributions of selection coefficients to each bin, and computed the mean divided by the standard deviation of each distribution, which is indicative of its shape (see Results). E) To test whether the gamma distributions were a significantly better fit than the single-coefficient mapping, we computed log-likelihood ratio statistics of the two models at each bin. The black line denotes the Bonferroni-corrected significance cutoff. F) To test how well we were fitting the data at each bin, we computed chi-squared statistics of the fitted SFS to the observed SFS at each bin. The black line denotes the Bonferroni-corrected significance cutoff.

We were concerned that our binning-based mapping would miss important features about the distribution of coefficients within each bin. To address this, we also fitted individual gamma distributions of selection coefficients to each of the bins. We show the mean, standard deviation (SD) and ancestral misidentification rate of each gamma fitting in Figure S3. The shape of the fitted gammas vary from an L-shape (Mean/SD < 1) at low C bins, to a shape resembling a skewed normal distribution at intermediate C bins (Mean/SD > 1) to a shape resembling an exponential distribution at high C bins (Mean/SD *≈* 1) (Figure 1.D). We performed a likelihood ratio test comparing the gamma model to the single-coefficient model, and find that only 4 out of the 40 bins (containing only 0.5% of all polymorphisms and 4.7% of nonsynonymous polymorphisms) are significantly supportive of the gamma model (Figure 1.E). A chi-squared test of the fit to the data shows both models perform similarly well, though both result in significant chi-squared scores at low C-score bins when using the genome-wide data (Figure 1.F). We attribute this to the large amount of data present in those bins as well as complex details of demographic history that affect neutral sites but that we did not model in our neutral fitting. In contrast, when mapping using only the exome, almost all bins have non-significant statistics, suggesting that selection dominates over demography in these regions. Based on these results, we decided to use the smoothed single-coefficient fitting for estimating the DFE in most downstream analyses, although we may be missing a small proportion of within-bin variability. Additionally, we show the inferred DFE of all, nonsynonymous and synonymous polymorphisms obtained from the gamma-fitted mapping in Figure S5.

We aimed to test the robustness of the selection coefficient estimates within each bin. We were specifically concerned about highly deleterious bins, which are composed of a smaller number of segregating sites than neutral or nearly neutral bins, and could produce unstable or biased estimates. We obtained bootstrapped confidence intervals for each bin and observe that the mappings are relatively stable up to C= 35 (Figure 1.A). As expected, the standard deviation of the bootstrap estimates is strongly negatively correlated with the sample-size per bin (Figure S6, Pearson correlation coefficient = −0.89). Thus, most of the increase in the width of the confidence intervals observed at higher C-score bins can be explained by the small number of polymorphisms available in those bins, and is likely not the result of other unaccounted processes, such as positive selection, operating exclusively on highly scored polymorphisms. To verify that our mapping was not an artifact of the different number of C-scores within each bin, we also performed 100 randomizations of the C-score assignments to each SNP in the genome. For each randomization, we re-computed the C-to-s mapping. When doing so, the bootstrap confidence intervals increase in size with increasing C scores, but the mapping remains flat, as expected (Figure 1.B).

Further, we verified that the mapping did not change considerably when filtering for sites in regions with low CpG density (*<* 0.05), defined as the proportion of CpG dinucleotides in a window of +/− 75 bp around the site [27] (Figure S7.A). This is expected, as the C-score model accounts for differential mutation rates at CpG sites and the resulting scores are generally robust to them [27]. As before, the gamma model is a significantly better fit than the single-coefficient model at only 4 out of the 40 bins (Figure S7.B).

Additionally, we re-mapped the scores using a constant-size model, and verified that the mapping does not change considerably if we assume a different demographic history than the best fit (Figure S8). The mappings are highly similar in shape, with the exception that, because the constant-size model is depleted of singletons relative to the best-fit model, it takes more bins to reach an SFS that is deleterious enough to map to *s* ≠ 0, and so more C-scores map to s= 0.

To test the dependence of our mapping on the choice of score used to estimate selection coefficients, we performed the same fitting procedure on a variety of other conservation and deleteriousness scores (see Methods). We note, however, that all of these scores are included as input in the C-score SVM. Figure S9 shows that the shape of the mapping is fairly consistent across different choices of scores, except for highly deleterious bins, which contain very few sites. When comparing different categories of sites in the Results, we show their distribution of selection coefficients obtained from the C-score mapping, as this score has been shown to be a better correlate to functional disruption than all the other scores mentioned above, and also controls for mutation rate variation across the genome, while other scores do not [27]. Additionally, we plot the mapped density of selection coefficients (with smoothing bandwidth = 0.000005) for all polymorphisms as well as synonymous and nonsynonymous polymorphisms using each of the other scores in Figure S10. We observe that, while all scores easily distinguish genic sites, PhastCons scores have difficulty distinguishing between synonymous and nonsynonymous sites, which we believe is due to PhastCons scores being regional, rather than position-specific scores. Additionally, while bimodality at genic sites is most prominent when using C-scores, it also becomes apparent in other position-specific scores when plotting the density with a finer smoothing bandwidth (= 0.000001, Figure S11).

### The distribution of fitness effects of segregating mutations

Using the C-score-to-selection coefficient mapping, we obtained the DFE of segregating polymorphisms in Yoruba individuals. This distribution is highly peaked when all polymorphisms are considered (Figure 2, black dashed line), with a large proportion of neutral changes and a long tail of deleterious mutations, as has been observed before when estimating the DFE of coding sequences [9, 11–13, 20]. Interestingly, we observe a pronounced drop in frequency for values of *s < −*10*^−^*^4^. We note that this is not due to our capping our mapping at *C*= 40 as the selection coefficients we are able to map are of a greater magnitude than this drop (Figure 1, S12).

**Figure 2.**
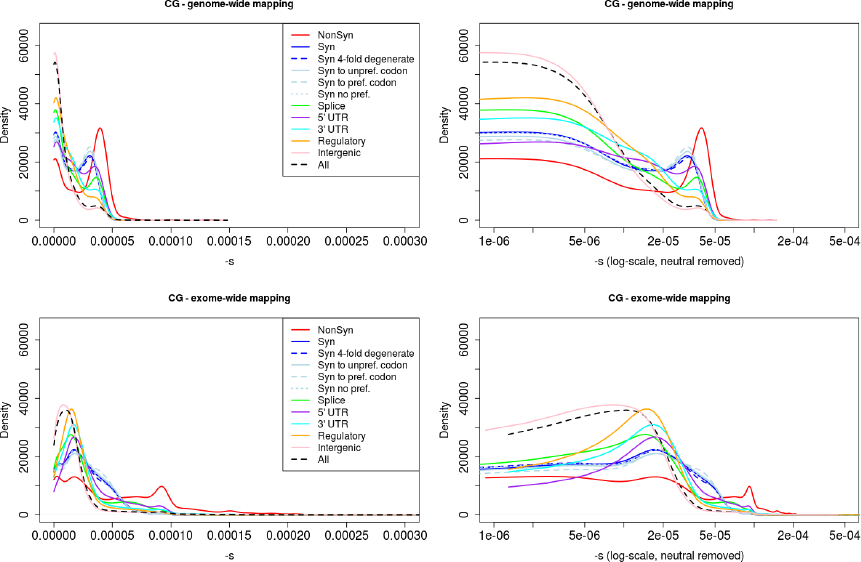
Distribution of fitness effects among YRI polymorphisms in the Complete Genomics dataset, partitioned by the genomic consequence of the mutated site. The right panels show a zoomed-in version of the same distributions after removing neutral polymorphisms and log-scaling the x-axis. Top row: DFE obtained from genome-wide mapping. Bottom row: DFE obtained from exome-wide mapping. Consequences were determined using the Ensembl Variant Effect Predictor (v.2.5). If more than one consequence existed for a given SNP, that SNP was assigned to the most severe of the predicted categories, following the VEP’s hierarchy of consequences. NonSyn = nonsynonymous. Syn = synonymous. Syn to unpref. codon = synonymous change from a preferred to an unpreferred codon. Syn to pref. codon = synonymous change from an unpreferred to a preferred codon. Syn no pref. = synonymous change from an unpreferred codon to a codon that is also unpreferred. Splice = splice site. CG = Complete Genomics data.

We partition the data by the genomic consequence of the polymorphisms, using the Ensembl Variant Effect Predictor (v.2.5) [36]. Some classes exhibit a peak of highly deleterious changes around *s*= *−*10*^−^*^4^. This peak results in a bimodal distribution that is especially pronounced for nonsynonymous sites (Figure 2, red line), and is almost non-existent for intergenic sites (Figure 2, pink line). Other types of polymorphisms—like splice site, synonymous, 3’ UTR, 5’ UTR and regulatory mutations—have less deleterious peaks than the one observed at nonsynonymous polymorphisms (Figure 2). We can compare the selection coefficient distributions to the distributions of unmapped C-scores (Figure S12) which are much less tightly peaked at intermediate C-score values and do not show a sharp decrease in density for high values, as does the s distribution in Figure 2. We show various statistics calculated on each of the selection coefficient distributions in Table 1 with the genome-wide mapping and in Table S1 with the exome-wide mapping.

**Table 1.** Characteristics of fitness effect estimated for YRI SNPs classified by different genomic consequence, RegulomeDB and Segway categories, using the genome-wide C-to-s mapping. We show quantiles of selection coefficients, the log base 10 of the mean selection coefficient and the log base 10 of the standard deviation of coefficients in each category.

| Category | n | $\vert s\vert < 10^{-5}$ | $\vert s\vert \geq 10^{-5}$ | $\vert s\vert \geq 5*10^{-5}$ | $\vert s\vert \geq 10^{-4}$ | $\log_{10}(\overline{-s})$ | $\log_{10}(SD(-s))$ |
| --- | --- | --- | --- | --- | --- | --- | --- |
| All | 9065398 | 73.77% | 26.23% | 0.09% | 0% | -5.16 | -4.95 |
| Nonsynonymous | 32522 | 28.88% | 71.12% | 2.07% | 0.14% | -4.6 | -4.74 |
| Synonymous | 30630 | 42.08% | 57.92% | 0.01% | 0% | -4.81 | -4.87 |
| Synonymous 4-fold degenerate | 16895 | 41.93% | 58.07% | 0% | 0% | -4.81 | -4.87 |
| Synonymous to unpreferred codon | 12498 | 40.01% | 59.99% | 0.01% | 0.01% | -4.79 | -4.86 |
| Synonymous to preferred codon | 4574 | 38.43% | 61.57% | 0% | 0% | -4.78 | -4.87 |
| Synonymous no preference change | 11844 | 41.80% | 58.20% | 0.02% | 0% | -4.81 | -4.87 |
| Splice site | 10122 | 52.11% | 47.89% | 0.05% | 0% | -4.87 | -4.84 |
| 5' UTR | 23125 | 37.53% | 62.47% | 0.11% | 0.02% | -4.76 | -4.84 |
| 3' UTR | 81605 | 49.05% | 50.95% | 0.19% | 0.02% | -4.88 | -4.87 |
| Regulatory | 864769 | 58.72% | 41.28% | 0.02% | 0% | -4.99 | -4.92 |
| Intergenic | 3544204 | 77.69% | 22.31% | 0.16% | 0% | -5.22 | -4.96 |
| GWAS | 8693 | 65.71% | 34.29% | 0.18% | 0% | -5.04 | -4.91 |
| eQTL+TF binding+matched TF motif+matched DNase Footprint+DNase peak | 240 | 47.08% | 52.92% | 0% | 0% | -4.86 | -4.88 |
| eQTL+TF binding+any motif+DNase Footprint+DNase peak | 1965 | 51.30% | 48.70% | 0.10% | 0% | -4.91 | -4.9 |
| eQTL+TF binding+matched TF motif+DNase peak | 62 | 66.13% | 33.87% | 0% | 0% | -5.01 | -4.89 |
| eQTL+TF binding+any motif+DNase peak | 1263 | 58.04% | 41.96% | 0.08% | 0% | -4.97 | -4.9 |
| eQTL+TF binding+matched TF motif | 44 | 65.91% | 34.09% | 0% | 0% | -4.95 | -4.83 |
| eQTL+TF binding / DNase peak | 26610 | 63.87% | 36.13% | 0.06% | 0% | -5.03 | -4.92 |
| TF binding+matched TF motif+matched DNase Footprint+DNase peak | 12411 | 50.05% | 49.95% | 0.10% | 0% | -4.87 | -4.87 |
| TF binding+any motif+DNase Footprint+DNase peak | 120642 | 57.03% | 42.97% | 0.12% | 0% | -4.95 | -4.89 |
| TF binding+matched TF motif+DNase peak | 5355 | 65.45% | 34.55% | 0.06% | 0% | -5.05 | -4.92 |
| TF binding+any motif+DNase peak | 96905 | 63.14% | 36.86% | 0.16% | 0% | -5.02 | -4.9 |
| TF binding+matched TF motif | 4359 | 73.34% | 26.66% | 0.07% | 0% | -5.15 | -4.95 |
| TF binding+DNase peak | 418548 | 59.95% | 40.05% | 0.10% | 0% | -4.99 | -4.91 |
| TF binding or DNase peak | 1625195 | 69.18% | 30.82% | 0.14% | 0% | -5.09 | -4.93 |
| C0 - CTCF (strong) | 20813 | 57.22% | 42.78% | 0.02% | 0.01% | -4.98 | -4.94 |
| C1 - CTCF (weak) | 61946 | 65.88% | 34.12% | 0.05% | 0% | -5.08 | -4.96 |
| D - dead zone | 873933 | 81.44% | 18.56% | 0.06% | 0% | -5.3 | -5 |
| E/GM - enhancer/gene middle | 70593 | 59.27% | 40.73% | 0.06% | 0% | -4.99 | -4.91 |
| F0 - FAIRE only | 1177948 | 71.94% | 28.06% | 0.11% | 0% | -5.13 | -4.94 |
| F1- FAIRE only | 1481766 | 74.45% | 25.55% | 0.10% | 0% | -5.17 | -4.95 |
| GE0 - gene body (end) | 413390 | 67.56% | 32.44% | 0.09% | 0% | -5.06 | -4.91 |
| GE1 - gene body (end) | 163365 | 63.99% | 36.01% | 0.10% | 0% | -5.03 | -4.91 |
| GE2 - gene body (end) | 54261 | 69.52% | 30.48% | 0.13% | 0.01% | -5.1 | -4.93 |
| GM0 - gene body (middle) | 130000 | 64.24% | 35.76% | 0.07% | 0% | -5.04 | -4.93 |
| GM1 - gene body (middle) | 101426 | 63.21% | 36.79% | 0.07% | 0% | -5.04 | -4.94 |
| GS - gene body (start) | 54940 | 41.21% | 58.79% | 0.06% | 0.01% | -4.83 | -4.89 |
| H3K9me1 | 457492 | 87.22% | 12.78% | 0.02% | 0% | -5.46 | -5.07 |
| L0 - low zone | 673274 | 81.71% | 18.29% | 0.07% | 0% | -5.31 | -5.01 |
| L1 - low zone | 725876 | 70.63% | 29.37% | 0.15% | 0% | -5.11 | -4.94 |
| N/A | 53354 | 95.36% | 4.64% | 0% | 0% | -5.77 | -5.29 |
| R0 - repression | 716710 | 74.73% | 25.27% | 0.12% | 0% | -5.17 | -4.95 |
| R1 - repression | 335824 | 69.73% | 30.27% | 0.09% | 0% | -5.1 | -4.94 |
| R2 - repression | 447655 | 68.85% | 31.15% | 0.13% | 0% | -5.08 | -4.91 |
| R3 - repression | 433156 | 75.62% | 24.38% | 0.09% | 0% | -5.19 | -4.97 |
| R4 - repression | 205971 | 73.26% | 26.74% | 0.06% | 0% | -5.15 | -4.94 |
| R5 - repression | 297152 | 70.84% | 29.16% | 0.09% | 0.01% | -5.13 | -4.96 |
| TF0 - transcription factor activity | 346083 | 73.50% | 26.50% | 0.09% | 0% | -5.17 | -4.97 |
| TF1 - transcription factor activity | 327695 | 77.18% | 22.82% | 0.06% | 0% | -5.22 | -4.98 |
| TF2 - transcription factor activity | 127366 | 71.09% | 28.91% | 0.12% | 0.01% | -5.12 | -4.95 |
| TSS - transcription start site | 25144 | 23.16% | 76.84% | 0.12% | 0.02% | -4.68 | -4.88 |

Next, we partitioned the data by whether the polymorphisms were found in the GWAS database [37] or not (Figure S13, Tables 1, S1). While we observe a second deleterious peak among the GWAS SNPs as well, these SNPs seem to be highly enriched for neutral polymorphisms.

Finally, we classified polymorphisms by different regulatory categories. We used two regulatory tracks downloaded from the UCSC Genome Browser [31, 38]. First, we partitioned the genome by regulatory regions identified by Regulome DB [39], which predicts regulatory activity in noncoding regions based on different types of experimental evidence (Figure S14, Tables 1, S1). Second, we used the Segway regulatory segment tracks [40], which are the product of an unsupervised pattern discovery algorithm that serves to identify and label regulatory regions along the genome, including genic regions (Figure 3, Tables 1, S1).

**Figure 3.**
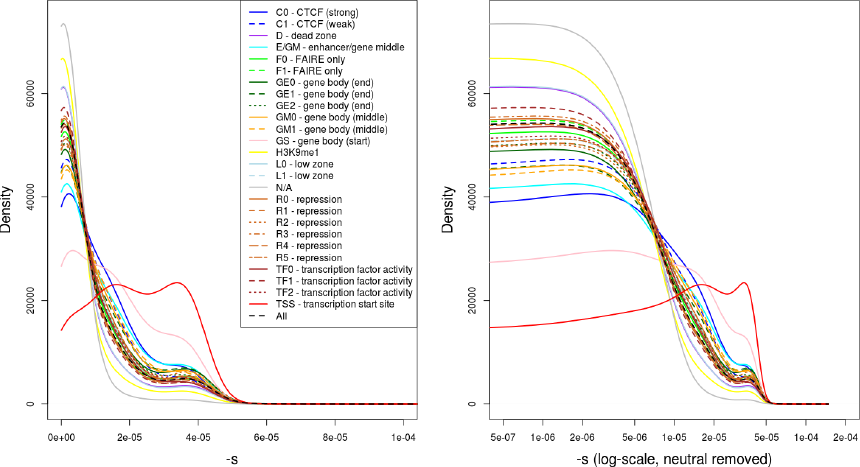
Distribution of fitness effects among YRI polymorphisms, partitioned by Segway regulatory segments. The right panel shows a zoomed-in version of the same distributions after removing neutral polymorphisms and log-scaling the x-axis.

## Discussion

The distribution of fitness effects (DFE) describes the proportion of mutations with given selection coefficients. Knowledge of the DFE has profound implications for our understanding of evolution and health. We infer a highly peaked distribution for all polymorphisms, with a drop in density at *s ≈ −*10*^−^*^4^, which may be the cutoff between weakly deleterious mutations that segregate in human populations and highly deleterious mutations that are easily pruned away by negative selection.

Our inferred non-synonymous distribution is bimodal and looks very similar to the one obtained for nonsynonymous mutations in Drosophila in ref. [11], with a peak at neutrality and another peak at *s ≈ −*10*^−^*^4^. Several experimental studies have also shown that non-synonymous non-lethal mutations tend to have a multimodal DFE in model organisms [41,42] (see ref. [18] for a comprehensive review). We note that it is impossible to obtain such kinds of distributions using a gamma or lognormal probability distribution unless one approximates bimodality by assuming a second, separate class of nonsynonymous mutations that are completely neutral and do not follow the best-fitting probability distribution [11, 13, 20, 25].

We also tested the precision of the C-scores by fitting gamma distributed DFEs to each C-score bin. While only very few bins were fit by a highly peaked gamma distribution (Figure 1D), the increased variation offered by the gamma distribution rarely improved the likelihood significantly (Figure 1E). This suggests that the C-scores are precise quantifications of negative selection, and that mutations with similar C-scores are likely to have similar selection coefficients.

Interestingly, we found that for many low C-score bins, a chi-squared test would reject the fit of the demographic model to the data. This is likely because these low C-score bins have a significant number of sites, and hence subtle features of the demography and quality control are relevant. Nonetheless, we note that for moderate-to-high C-score bins and for exonic data, we would not be able to reject the fit of the the predicted site frequency spectrum from the data. While these bins have fewer sites, it is also likely that stronger selection is obscuring some of the signal of subtle demographic events.

Our novel expectation-maximization approach to quantifying ancestral state misidentification allows us to assess differential misidentification rates across C-score categories. Ancestral state misidentification occurs because a site is hit by more than one mutation, hence obscuring the identity of the ancestral allele. Here, we found that low C-score bins are enriched with ancestral state misidentification, with rates in excess of 5% for some bins (Figure S3). This may reflect a bias induced by the C-scores themselves, because C-scores are trained to distinguish classes of sites that have relatively few substitutions between humans and chimpanzees. Because the signal of ancestral state misidentification and positive selection are very similar [43], it is possible that low C-score bins are enriched for positive selection, although we did not pursue that direction any further. For larger C-score bins, we infer misidentification rates along 10 the lines of those obtained in simulation studies by ref. [43].

Importantly, unlike previous studies, we also obtain DFEs for other types of mutations, including synonymous, splice site, 3’ UTR, 5’ UTR and regulatory polymorphisms, which exhibit bimodality to a lesser degree than the nonsynonymous DFE. In particular, 5’ UTR changes constitute the category with the smallest proportion of neutral or nearly neutral (*|s| <* 10^−5^) polymorphisms after nonsynonymous changes, likely reflecting selection on gene regulation upstream of coding sequences. Futhermore, distributions corresponding to mutations in UTR and regulatory regions have a less pronounced trough between the two peaks than the ones observed among coding mutations, suggesting that the magnitude of deleterious effects is more uniformly distributed in non-coding regions. In contrast, missense mutations appear to have more of an “all-or-nothing” effect, as would perhaps be expected when replacing an amino acid inside a protein.

Our method does not use synonymous sites as a neutral benchmark, as do other studies [9, 11, 20]. In fact, our inferred DFE suggests that a considerable number of synonymous polymorphisms may not be neutral, as we observe a second deleterious peak in them too, albeit less deleterious than the one observed at nonsynonymous polymorphisms. We caution, however, that this second peak is less promient when using an exome-specific mapping (Figure 2) or when using strictly position-specific scores (Figures S10, S11), which suggests that at least part of this peak may be caused by regional patterns of conservation or background selection in the exomes. Instead, intergenic polymorphisms are the class of sites most likely to evolve neutrally. Because this class is so abundant, most of the signal observed when all polymorphisms are pooled together closely reflects the distribution observed for intergenic polymorphisms (Figure 2).

Our results have implications for GWAS, as we find a high proportion of GWAS SNPs to be neutral or nearly neutral, which could suggest a high rate of false positives in this type of association studies. On the other hand, GWAS studies only aim to find polymorphisms linked to causative variants, and so GWAS SNPs need not have strongly deleterious effects. Alternatively, if the effect size of many GWAS SNPs are sufficiently small, it is possible that many of them are not subject to strong selection.

Additionally, by stratifying our results based on different ENCODE categories, we can elucidate the fitness consequences of molecular activity signals detected by ENCODE [2, 3, 39]. We find the category with the lowest proportion of neutral polymorphisms to be the one corresponding to sites that have eQTL evidence as well as evidence for transcription factor (TF) binding, a matched TF motif, a matched DNase footprint and that are located in a DNase peak. In general, categories that combine many regulatory 11 signals tend to show lower proportions of neutral mutations than those that do not, suggesting that data integration across distinct approaches to detecting selection and functionality is likely to do better than any individual approach [44]. Moreover, this suggests that much of the molecular activity detected by ENCODE may not have significant fitness consequences.

Stratification by Segway regions allows us to look at a different aspect of regulatory activity in the genome. Interestingly, we observe that polymorphisms in Transcription Start Sites (TSS) are the ones containing the largest proportion of deleteriousness. This echoes results from analyses of variation at transcription factor binding sites, which have been found to be under stronger constraint when found near TSS than when found far from them [45]. Other regions with high proportions of deleterious polymorphisms include Gene Body (Start), strong CTCF and Enhancer / Gene Middle. This suggests that regions with strong repressor, insulator or enhancer activity, as well as near the start of genes, are particularly important for biological function, perhaps unsurprisingly given our knowledge of the molecular role that these regions play in the regulation of transcription.

There are several limitations to our method. First, we have restricted ourselves to estimating the DFE of segregating mutations that have reached appreciable frequencies in the population. An extension of this approach would be to infer the DFE of new mutations from the DFE of segregating mutations genome-wide. Second, we assumed no dominance, epistasis or positive selection, which future studies could attempt to incorporate into our approach. In addition, we have assumed sites are independent and have therefore ignored the covariance between linked sites, which likely leads to an underestimatation of confidence intervals obtained from the bootstrapping. The free-recombination assumption may also affect inference due to Hill-Robertson interference between mutations subject to selection [46] as well as linked background selection affecting the SFS of neutral sites in the human genome [35]. This may be a more important issue in our case than other genic-only approaches because we are also including intergenic mutations in our analysis, so the space between analyzed polymorphisms is on average smaller than if we were only looking at coding polymorphisms [20]. We also assume no positive selection. This, however, should not be a major problem, because we are only basing our inferences on polymorphic sites and advantageous mutations contribute little to polymorphism, assuming *N_e_s >* 25 [32]. One final limitation is that the type of inference performed here is only possible in species in which C-scores have been estimated (for now, humans only) and that it therefore relies on C-scores being able to correctly rank sites by deleteriousness. Nevertheless, it should not be hard to obtain accurate C-scores for other 12 organisms in the future, although limitations on available annotations for non-human organisms may make the approximation to the fitness distribution less accurate than in humans.

## Materials and Methods

### Computing the theoretical site frequency spectrum

We used the theory developed by Evans *et al.* [47] to obtain the expected population site frequency spectrum with non-equilibrium demography. We assume a Wright-Fisher population in the limit of large population size and use diffusion theory to model this process. Writing *f* (*x, t*) for the frequency spectrum at frequency x and time *t* = 2*N* (0)*τ* where *τ* is in units of generations and *g*(*x, t*):= *x*(1 −*x*)*f* (*x, t*), we can approximate the dynamics of *g*(*x, t*) with genic selection and mutation by solving the following partial differential equation:

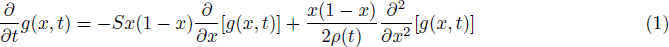

subject to the boundary condition:

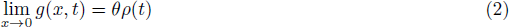

where S is the population-scaled selection coefficient (*S* = 2*N* (0)*s*), *θ* is the population-scaled mutation rate (*θ*= 4*N* (0)*µ*) and *ρ*(*t*) = *N* (*t*)*/N* (0) is the effective population size at time *t* relative to the population size at time 0.

We assume N(0) to be 10,000 for all exponential and constant models. For the constant population size model, *ρ*(*t*) = 1. For the exponential growth model *ρ*(*t*) = *exp*(*rt*) where *r* = 2*N* (0)*R* is the population-scaled growth rate and *R* is the per-generation growth rate. For models taken from the literature, we use the N(0) assumed by the corresponding paper. For the model of Harris and Nielsen, *ρ*(*t*) is piece-wise defined according to their Figure 7. The Tennessen model is similarly defined in a piece-wise fashion according to their Figure 2, although it also includes periods of exponential growth, as opposed to simply being piece-wise constant as in the Harris and Nielsen model.

We solve for *g*(*x, t*) numerically in Mathematica, and can then compute the expected number of segregating sites with *i* copies of the derived allele out of a sample of *n* genes. Denoting by *g_s_*(*x, t*) the theoretical SFS with selection coefficient *s*, this quantity is

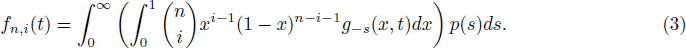

where *p*(*s*) is the parameterized distribution of selection coefficients. For gamma distributed fits,

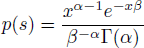

where *α* and *β* are the shape and rate parameters of the gamma distribution and Γ(*·*) is the gamma function. For a point mass at 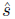,

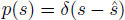

where *δ*(*·*) is the usual Dirac delta function.

We focused on fitting the shape of the SFS, and hence worked with the probability that a given site in a sample of *n* has *i* copies of the derived allele,

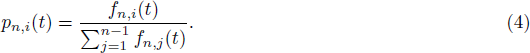

The Mathematica code used for obtaining the theoretical SFS can be found online at: http://malecot.popgen.dk/schraiber/

### Expectation maximization algorithm to fit ancestral state misidentification rates

We observed that the SFS showed signs of ancestral state misidentification, in particular an excess of high frequency derived alleles (Figure S2). To account for the ancestral state misidentification errors, we developed an expectation maximization (EM) algorithm. In the E step, we estimate the posterior probability that a site at frequency *i* out of *n* is misidentified given the current estimated site frequencies, 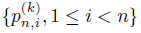, and the current estimate of the misidentification rate, 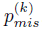, as

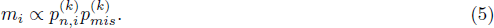

Then, during the M step, we re-estimate the misidentification rate as

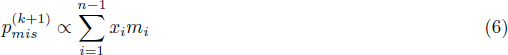

where *x_i_* is the number of sites at frequency *i*. We next re-estimate either the demographic parameters or the parameters of the selected model using the log-likelihood

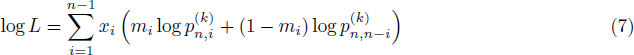

to obtain the theoretical SFS for the next iteration, 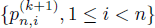

### Maximum likelihood fitting of demographic models

The exponential growth model has two free parameters, *r*, the population-scaled growth rate and *t*, the total time of exponential growth. We first obtained the site frequency spectrum for all sites with *C* = 0. Next we solved *g*(*x, t*) for the exponential growth model across a grid of *t* and *r*, as well as the constant, Harris and Tennessen models, and applied our EM algorithm to estimate the best fitting demographic model, using a grid search over demographic models during the M step.

### Maximum likelihood fitting of selection coefficients

To find the maximum likelihood estimate of *s* for each C-score bin, we first obtained the site frequency spectrum corresponding to each C-score bin. Next, we solved *g*(*x, t*) under the fitted demography for log_10_(*−s*) ∈ [−7*, −*1.3] in steps of 0.005, along with *s* = 0. To obtain an estimated SFS under the assumption of gamma distributed selection coefficients, we used the trapezoid rule to numerically integrate against a gamma distribution as in formula 3.

We used our EM algorithm to estimate the best fitting selection coefficient for each bin. When fitting a single coefficient, we used a grid search during the M-step, and when fitting gamma distributed selection coefficients, we used the L-BFGS-B algorithm. To plot the DFE, we used kernel density estimation with smoothing bandwith = 0.000005, unless otherwise stated.

### Mapping using different scores

To test how robust the mapping of C-scores to selection coefficients is to different types of conservation scores, we obtained PhyloP [48] and PhastCons [49] scores derived from vertebrate, mammal and primate alignments (excluding humans), as well as GERP++ rejected substitution (Gerp S) scores [50], for all YRI SNPs [27]. We attempted to equalize the range of all scores by PHRED-scaling them, i.e. converting each score to −log_10_(*p*) where *p* is the probability of observing a change as or more disruptive / conserved (based on that particular score scale) among all polymorphic YRI sites. We note that this is different from the natural PHRED scale of C-scores (where *p* is the the probability of observing a score as or more disruptive among all possible, but not necessarily realized, mutations in the human genome), and so we also re-scaled the C-scores to produce a fair comparison. Then, we repeated the maximum likelihood mapping for each PHRED-scaled score in bins of 0.25 units (e.g. 0-0.125, 0.125-0.375, 0.375-0.625, etc). It is important to note that PhastCons are regional scores, while PhyloP and Gerp S are position-specific scores. Another difference is that PhastCons scores only measure the probability of negative selection, while PhyloP and Gerp S scores also measure positive selection (i.e. low/negative scores represent faster evolution than expected purely under drift), which may be why we observe an uptick at the lower end of the mapping for those scores in Figure S9.

## Acknowledgments

We thank Montgomery Slatkin and Rasmus Nielsen for helpful advice and discussions, and Martin Kircher for providing the C-score data and other annotations. This work was supported by the National Institutes of Health (R01-GM40282 grant to Montgomery Slatkin).

## Supplementary Figures

**Figure S1.**
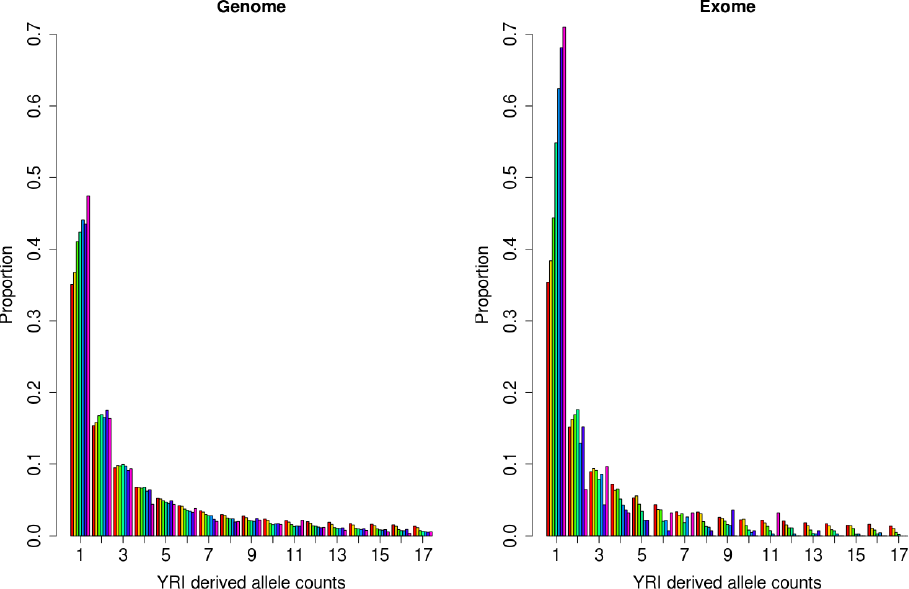
Observed SFS for sites under different C-score bins using the Complete Genomics YRI data, for all autosomes in the genome (left) and the exome (right). Note that the spectrum gets more skewed towards singletons with increasing C-scores, likely reflecting the action of negative selection on deleterious mutations.

**Figure S2.**
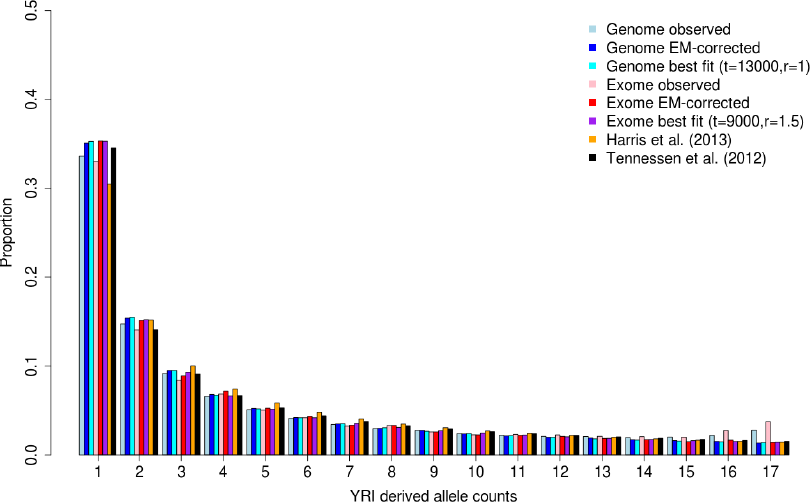
Observed SFS of YRI Complete Genomics data for sites with C = 0. The full SFS was corrected for ancestral state misidentification using an EM algorithm and fit to different models of neutral evolution. We show results for both the genome and the exome.

**Figure S3.**
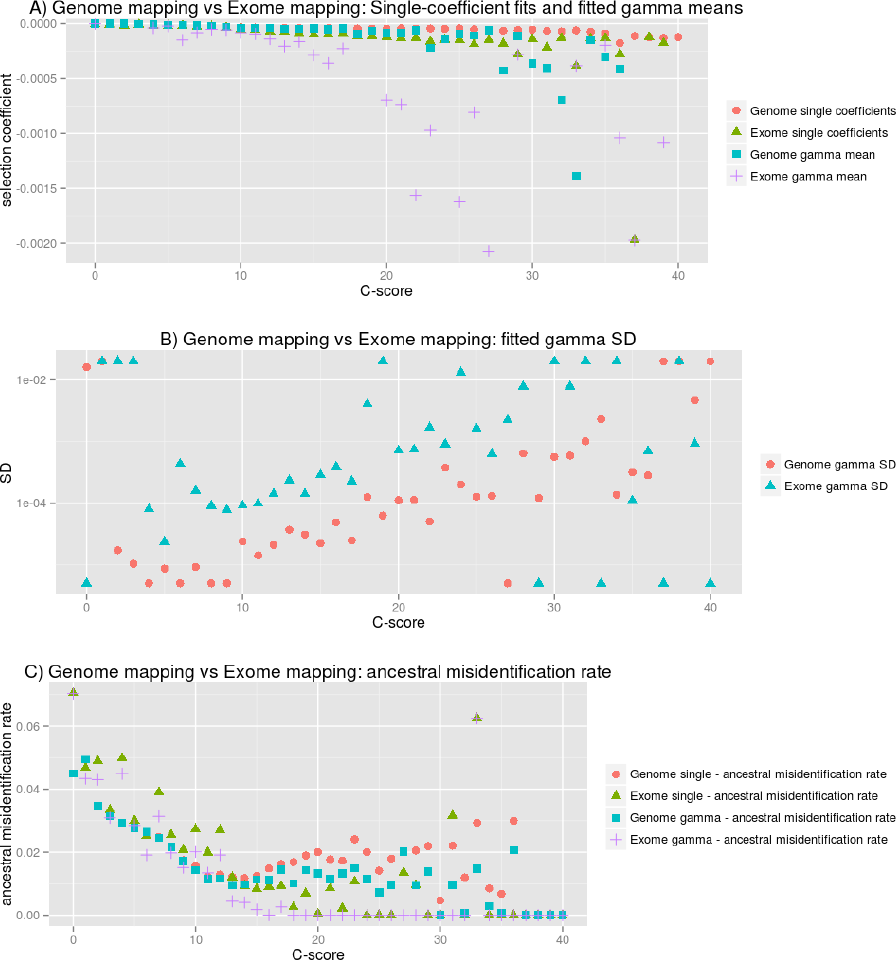
Features of fitted single-coefficient and gamma distributions. A) Fitted single coefficients and means of fitted gamma distributions for each C-score bin, using genome-wide or exome-wide polymorphisms. B) Standard deviation of fitted gamma distributions for each bin. C) Ancestral misidentification rate obtained from an EM algorithm used to jointly fit the data and infer this rate at each bin. SD = standard deviation.

**Figure S4.**
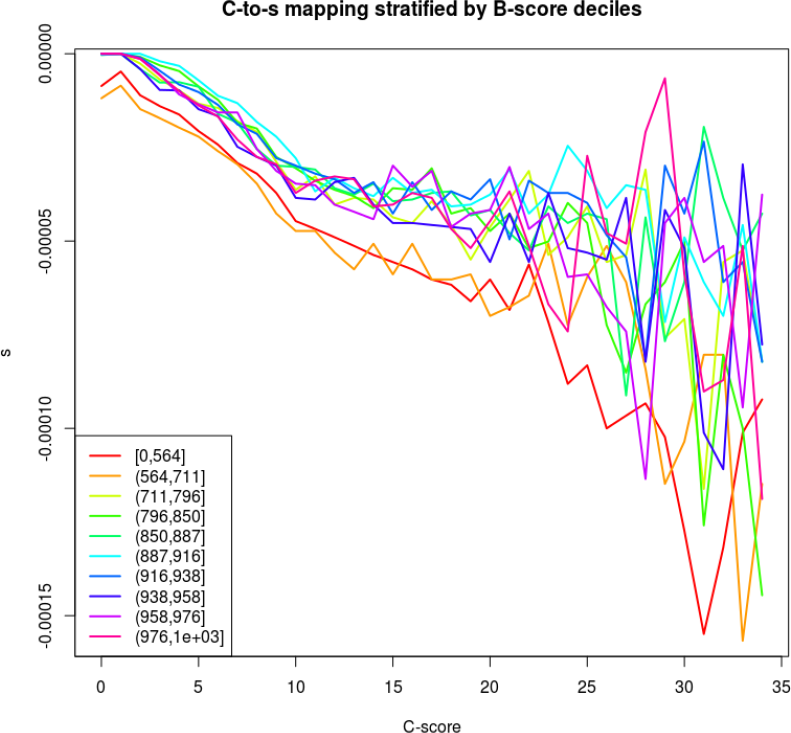
C-to-s mapping stratified by B-score deciles. We partitioned the genome by deciles of B-scores [35], which reflect levels of background selection. Then, we recomputed the demographic fitting and C-to-s mapping for each decile.

**Figure S5.**
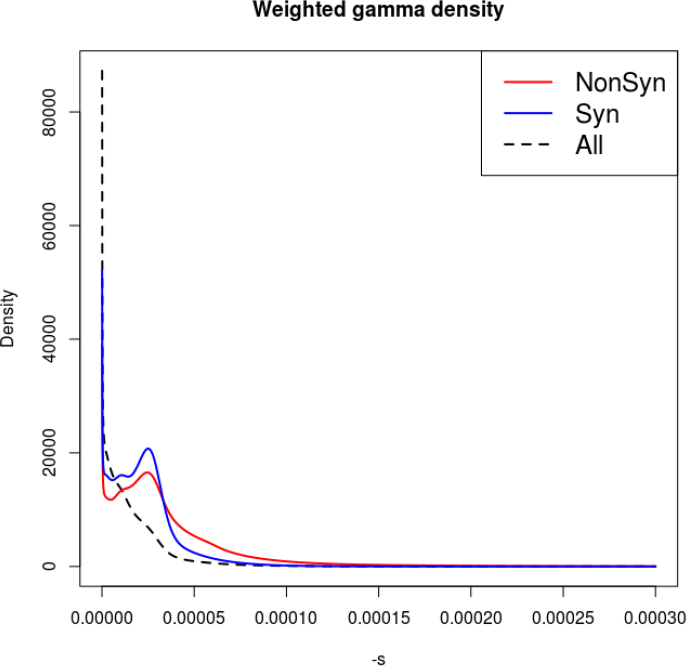
Inferred DFEs for all, nonsynonymous and synonymous polymorphisms obtained from gamma distribution fitting of each C-score bin. The plot shows, for each category, a weighted sum of gamma distributions, where each C-score bin contributes its corresponding genome-wide best-fitting gamma distribution in proportion to the number of polymorphisms present at that bin.

**Figure S6.**
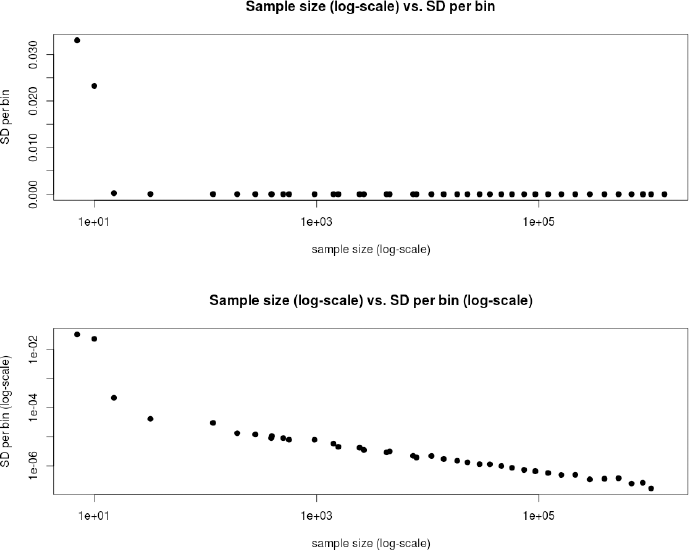
Comparison between the size of each C-score bin and the standard deviation of single-coefficient fits obtained from 100 bootstraps of the data within each bin. Top panel: Standard deviation per C-score bin plotted as a function of sample size per bin (log-scale). Bottom panel: Same plot but with the y-axis on a log-scale.

**Figure S7.**
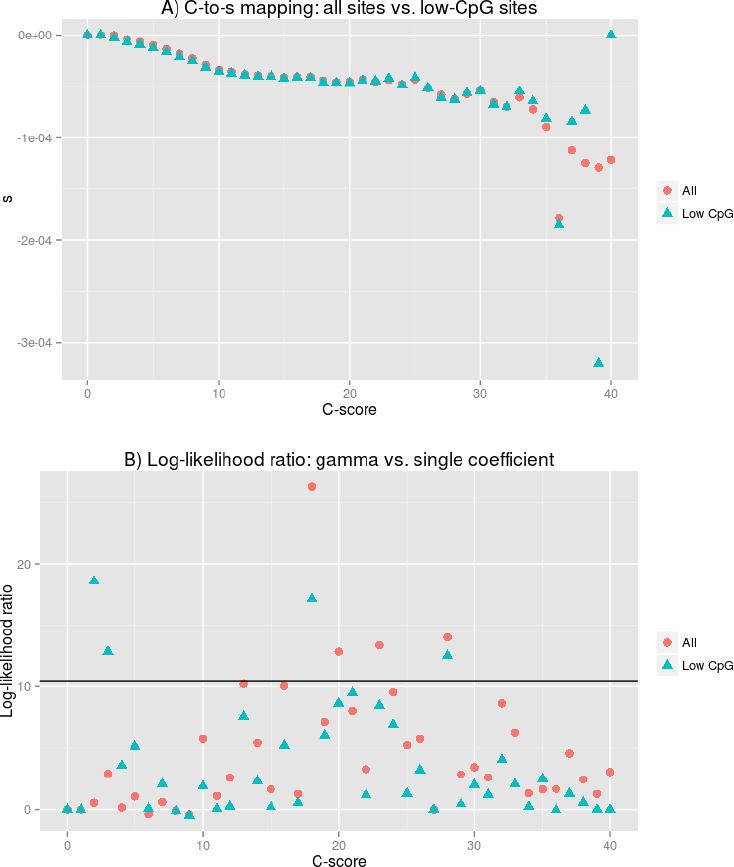
Mapping of sites with low CpG density. A) We filtered for sites with low CpG density, such that the proportion of CpG sites in a +/− 75 bp window around each site was *<* 0.05, and then recomputed the C-to-s mapping. B) We also repeated the gamma fitting and calculated a liklelihood ratio test of the gamma model against the single-coefficient model at each C-score bin.

**Figure S8.**
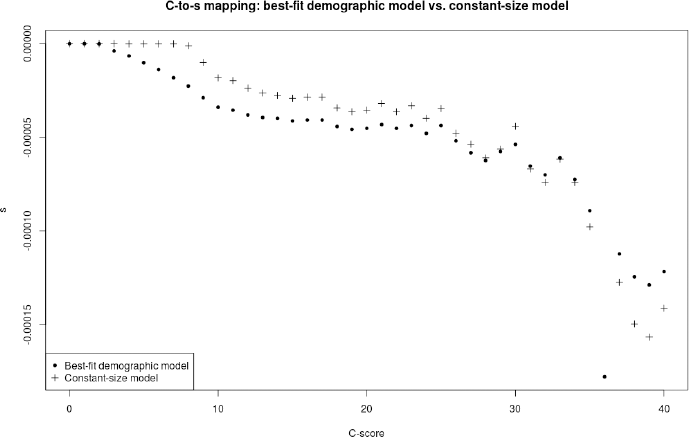
Comparison between a C-to-s mapping using the best-fit demographic model (exponential growth with t = 13,000 and r = 1) and a constant-size model.

**Figure S9.**
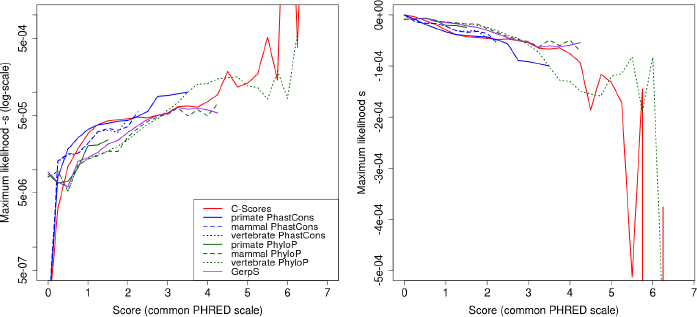
Maximum likelihood mapping of different types of scores to a selection coefficient scale, excluding bins mapped to neutrality, using the Complete Genomics data. Before mapping, scores were re-scaled on a common PHRED scale, by converting each score to −log_10_(*p*) where *p* is the probability of observing a change as or more disruptive / conserved (based on that particular score scale) among all polymorphic YRI sites. Some scores extend over larger PHRED scores than others because they have a finer stratification (smaller number of sites with tied scores). The wide fluctuations to the right of the figures are due to the small number of sites per bin at highly deleterious bins.

**Figure S10.**
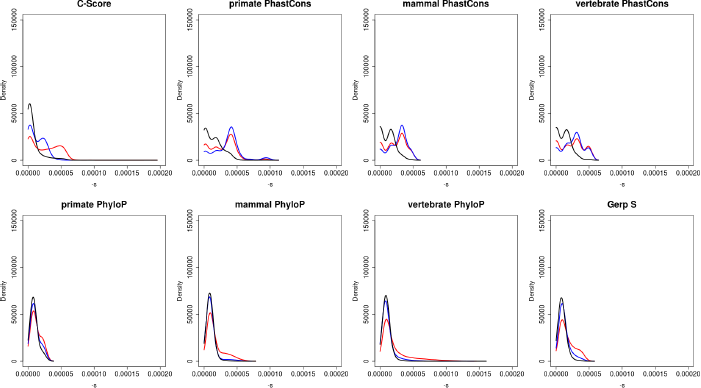
Distribution of fitness effects at nonsynonymous (red), synonymous (blue) and all (black) polymorphisms in Yoruba, using different types of conservation scores for mapping (smoothing bandwidth = 0.000005). We only mapped sites with PHRED-scaled scores ≤ 5, because the mappings become erratic for higher values, due to the small number of sites per bin (Figure S9).

**Figure S11.**
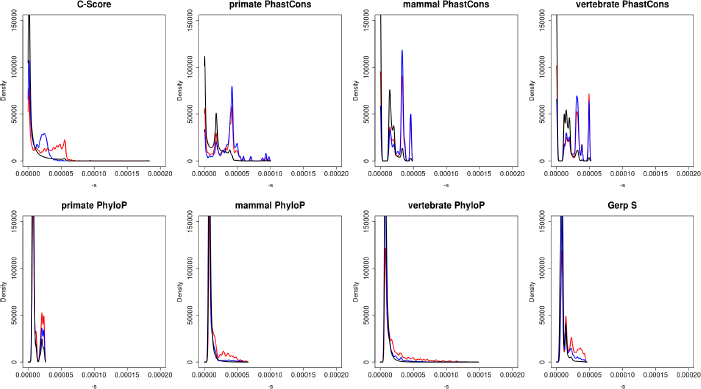
Distribution of fitness effects at nonsynonymous (red), synonymous (blue) and all (black) polymorphisms in Yoruba, using different types of conservation scores for mapping (smoothing bandwidth = 0.000001). We only mapped sites with PHRED-scaled scores ≤ 5, because the mappings become erratic for higher values, due to the small number of sites per bin (Figure S9).

**Figure S12.**
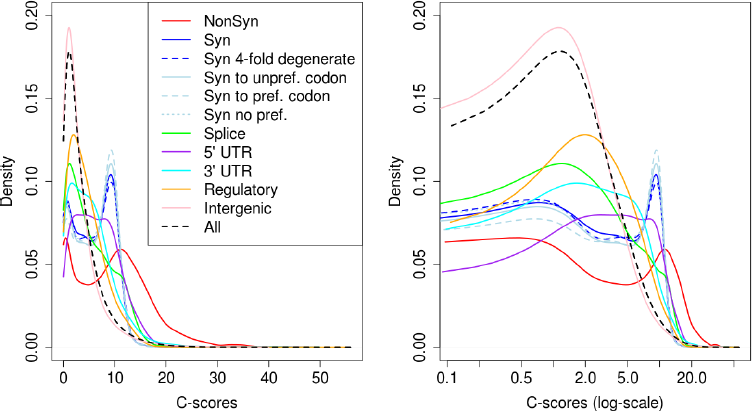
Distribution of unmapped C-scores among YRI polymorphisms, partitioned by the genomic consequence of the mutated site. Consequences were determined using the Ensembl Variant Effect Predictor (v.2.5). NonSyn = nonsynonymous. Syn = synonymous. Syn to unpref. codon = synonymous change from a preferred to an unpreferred codon. Syn to pref. codon = synonymous change from an unpreferred to a preferred codon. Syn no pref. = synonymous change from an unpreferred codon to a codon that is also unpreferred. Splice = splice site.

**Figure S13.**
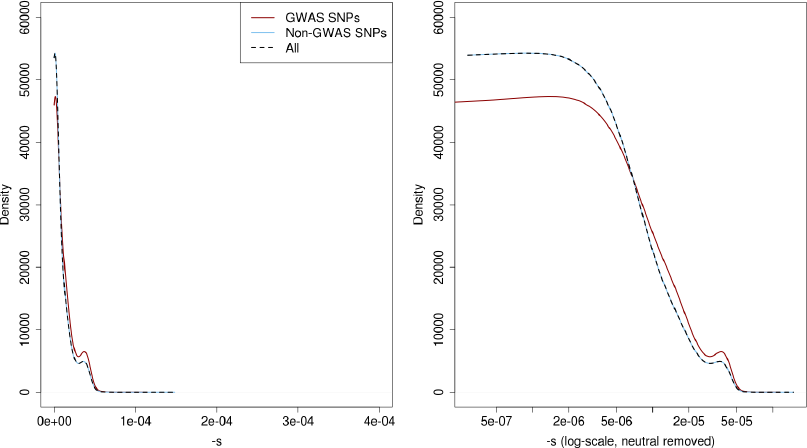
Distribution of fitness effects among YRI polymorphisms, partitioned by whether the SNPs are found in the GWAS database or not. The right panel shows a zoomed-in version of the same distributions after removing neutral polymorphisms and log-scaling the x-axis.

**Figure S14.**
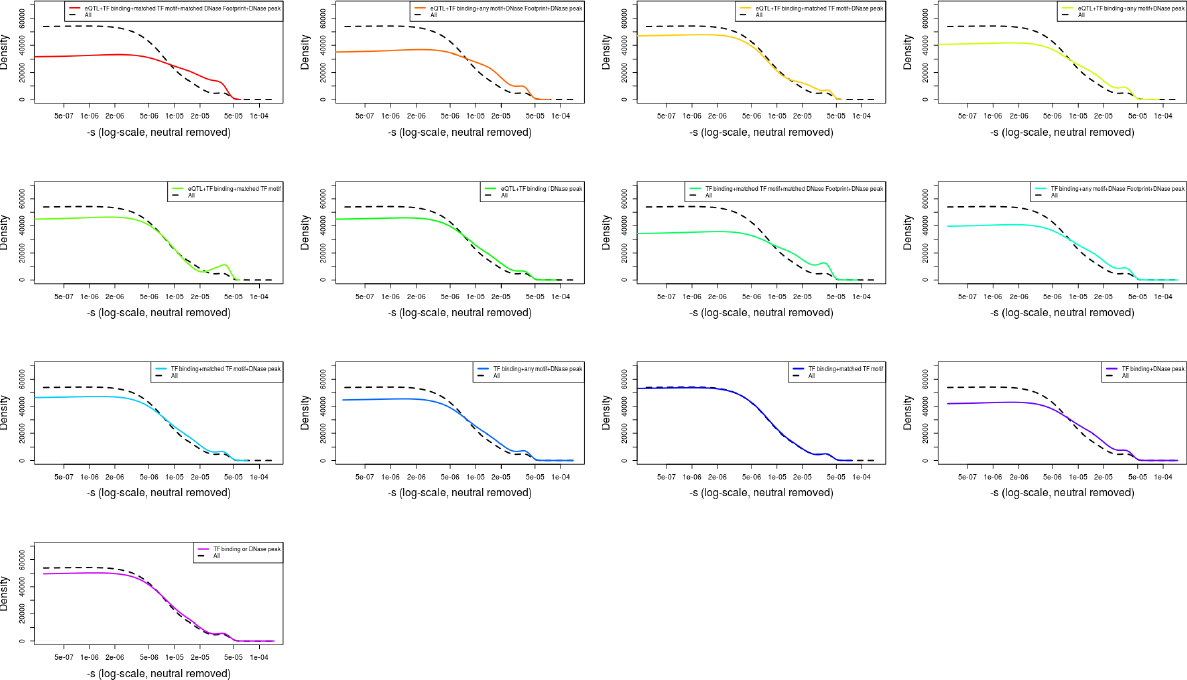
Distribution of fitness effects among different types of RegulomeDB regulatory YRI polymorphisms, obtained from various ENCODE assays. The black dashed line corresponds to the distribution of all YRI SNPs.

## Supplementary Tables

**Table S1.** Characteristics of fitness effect distributions estimated for YRI SNPs classified by different genomic consequence categories, RegulomeDB categories and Segway categories, using the exome-wide C-to-s mapping. We show quantiles of selection coefficients, the log base 10 of the mean selection coefficient and the log base 10 of the standard deviation of coefficients in each category.

| Category | n | $\vert s\vert < 10^{-5}$ | $\vert s\vert \geq 10^{-5}$ | $\vert s\vert \geq 5*10^{-5}$ | $\vert s\vert \geq 10^{-4}$ | $\log_{10}(\overline{-s})$ | $\log_{10}(SD(-s))$ |
| --- | --- | --- | --- | --- | --- | --- | --- |
| All | 9065398 | 40.51% | 59.49% | 5.87% | 0.92% | -4.79 | -4.7 |
| Nonsynonymous | 32522 | 17.26% | 82.74% | 45.87% | 11.52% | -4.28 | -4.35 |
| Synonymous | 30630 | 22% | 78% | 9.77% | 0.06% | -4.61 | -4.74 |
| Synonymous 4-fold degenerate | 16895 | 22.79% | 77.21% | 10.70% | 0.03% | -4.61 | -4.73 |
| Synonymous to unpreferred codon | 12498 | 21.54% | 78.46% | 10.52% | 0.10% | -4.59 | -4.73 |
| Synonymous to preferred codon | 4574 | 19.52% | 80.48% | 10.08% | 0% | -4.59 | -4.75 |
| Synonymous no preference change | 11844 | 22.55% | 77.45% | 9.58% | 0.03% | -4.61 | -4.74 |
| Splice site | 10122 | 25.43% | 74.57% | 16.08% | 0.42% | -4.6 | -4.63 |
| 5' UTR | 23125 | 13.95% | 86.05% | 18.65% | 0.88% | -4.52 | -4.55 |
| 3' UTR | 81605 | 20.98% | 79.02% | 11.94% | 1.60% | -4.6 | -4.58 |
| Regulatory | 864769 | 23.20% | 76.80% | 7.42% | 0.14% | -4.7 | -4.75 |
| Intergenic | 3544204 | 44.21% | 55.79% | 5.35% | 1.12% | -4.82 | -4.69 |
| GWAS | 8693 | 30.16% | 69.84% | 8.24% | 1.69% | -4.7 | -4.64 |
| eQTL+TF binding+matched TF motif+matched DNase Footprint+DNase peak | 240 | 14.58% | 85.42% | 11.67% | 0.83% | -4.6 | -4.7 |
| eQTL+TF binding+any motif+DNase Footprint+DNase peak | 1965 | 18.78% | 81.22% | 9.97% | 0.51% | -4.64 | -4.69 |
| eQTL+TF binding+matched TF motif+DNase peak | 62 | 25.81% | 74.19% | 8.06% | 1.61% | -4.69 | -4.65 |
| eQTL+TF binding+any motif+DNase peak | 1263 | 22.33% | 77.67% | 9.11% | 0.55% | -4.67 | -4.7 |
| eQTL+TF binding+matched TF motif | 44 | 31.82% | 68.18% | 13.64% | 2.27% | -4.62 | -4.56 |
| eQTL+TF binding / DNase peak | 26610 | 27.75% | 72.25% | 7.16% | 0.86% | -4.71 | -4.7 |
| TF binding+matched TF motif+matched DNase Footprint+DNase peak | 12411 | 17.94% | 82.06% | 13.42% | 0.54% | -4.6 | -4.65 |
| TF binding+any motif+DNase Footprint+DNase peak | 120642 | 22.76% | 77.24% | 9.62% | 0.82% | -4.66 | -4.66 |
| TF binding+matched TF motif+DNase peak | 5355 | 30.51% | 69.49% | 7.79% | 1.05% | -4.72 | -4.68 |
| TF binding+any motif+DNase peak | 96905 | 28.49% | 71.51% | 8.61% | 1.29% | -4.69 | -4.64 |
| TF binding+matched TF motif | 4359 | 38.63% | 61.37% | 6.33% | 0.99% | -4.78 | -4.7 |
| TF binding+DNase peak | 418548 | 24.64% | 75.36% | 8.15% | 0.87% | -4.68 | -4.68 |
| TF binding or DNase peak | 1625195 | 33.43% | 66.57% | 7.02% | 1.32% | -4.74 | -4.66 |
| C0 - CTCF (strong) | 20813 | 18.32% | 81.68% | 6.75% | 0.15% | -4.69 | -4.77 |
| C1 - CTCF (weak) | 61946 | 27.65% | 72.35% | 5.21% | 0.46% | -4.75 | -4.77 |
| D - dead zone | 873933 | 52.91% | 47.09% | 4.24% | 0.65% | -4.89 | -4.75 |
| E/GM - enhancer/gene middle | 70593 | 23.64% | 76.36% | 7.94% | 0.67% | -4.68 | -4.7 |
| F0 - FAIRE only | 1177948 | 36.88% | 63.12% | 6.41% | 1.04% | -4.76 | -4.69 |
| F1- FAIRE only | 1481766 | 41.02% | 58.98% | 5.94% | 1.03% | -4.79 | -4.69 |
| GE0 - gene body (end) | 413390 | 33.96% | 66.04% | 8.03% | 1.09% | -4.72 | -4.66 |
| GE1 - gene body (end) | 163365 | 28.37% | 71.63% | 8.06% | 1.03% | -4.7 | -4.67 |
| GE2 - gene body (end) | 54261 | 32.46% | 67.54% | 6.38% | 0.94% | -4.74 | -4.66 |
| GM0 - gene body (middle) | 130000 | 27.23% | 72.77% | 7.27% | 0.82% | -4.71 | -4.7 |
| GM1 - gene body (middle) | 101426 | 24.46% | 75.54% | 6.44% | 0.62% | -4.71 | -4.73 |
| GS - gene body (start) | 54940 | 10.49% | 89.51% | 11.35% | 0.45% | -4.58 | -4.7 |
| H3K9me1 | 457492 | 62.34% | 37.66% | 2.70% | 0.27% | -4.98 | -4.85 |
| L0 - low zone | 673274 | 52.14% | 47.86% | 4.02% | 0.67% | -4.89 | -4.76 |
| L1 - low zone | 725876 | 33.64% | 66.36% | 6.32% | 1.24% | -4.75 | -4.68 |
| N/A | 53354 | 35.18% | 64.82% | 0.87% | 0.07% | -4.96 | -5.09 |
| R0 - repression | 716710 | 42.48% | 57.52% | 6.18% | 1.11% | -4.79 | -4.68 |
| R1 - repression | 335824 | 32.41% | 67.59% | 6.33% | 1.01% | -4.75 | -4.69 |
| R2 - repression | 447655 | 34.45% | 65.55% | 7.83% | 1.37% | -4.73 | -4.65 |
| R3 - repression | 433156 | 41.40% | 58.60% | 5.10% | 0.85% | -4.81 | -4.72 |
| R4 - repression | 205971 | 41.10% | 58.90% | 6.35% | 0.80% | -4.78 | -4.7 |
| R5 - repression | 297152 | 31.70% | 68.30% | 5.15% | 0.66% | -4.77 | -4.74 |
| TF0 - transcription factor activity | 346083 | 35.10% | 64.90% | 4.81% | 0.69% | -4.79 | -4.75 |
| TF1 - transcription factor activity | 327695 | 45.07% | 54.93% | 4.85% | 0.65% | -4.83 | -4.74 |
| TF2 - transcription factor activity | 127366 | 33.95% | 66.05% | 5.69% | 0.82% | -4.76 | -4.68 |
| TSS - transcription start site | 25144 | 5.01% | 94.99% | 21.84% | 0.89% | -4.46 | -4.6 |

